# A Novel Open-Source CellProfiler Pipeline for Automated, User-Friendly Hierarchical and K-Means Clustering of Microglial Morphology

**DOI:** 10.64898/2026.09.24.753926

**Authors:** Antonia Landwehr, Cody Cools-Lartigue, Victor Hadera, Bianca Bobotis, Mohammadparsa Khakpour, Felipe V. Gomes, Marie-Ève Tremblay

## Abstract

Microglia represent a highly dynamic and heterogeneous cell type that is critically implicated in states of health and pathology. Microglial morphological subgroups have been identified that correspond to functional characteristics determining health-related outcomes. The identification of states based on morphological characteristics will therefore provide invaluable insights into the microglia-specific functional mechanisms driving treatment effects. The application of clustering analyses enables the detection of groupings within samples reflecting differences in morphological features. Here we propose the application of three custom-created modules to be used within the open-source software CellProfiler. These modules enable the automated detection of clusters present within the sample of microglia, as well as the assessment of the abundance of these clusters across conditions. The application of the analysis is conducted in a highly user-friendly manner, with a user interface integrated into the pipeline, enabling the performance of the analysis with only minimal user input. The workflow thereby includes the conduction of an outlier assessment, followed by hierarchical clustering and k-means clustering and the generation of interactive graphs to determine the number of microglia states present in the sample. Bar plots displaying the abundance of the microglia states across conditions included in the sample will be created. This approach will facilitate faster and more comparable detection of microglial morphological clusters across studies.

## 1. Introduction

Microglia, the macrophages of the central nervous system (CNS), are a highly dynamic type of cell, continuously adjusting their morphology in response to even minor changes in their microenvironment (A et al., 2005). Microglia fulfill a multitude of functions across health and pathology, being involved, for example, in the shaping of neuronal circuits, adjustment of the vasculature, and neurogenesis (Bisht et al., 2021; De Lucia et al., 2016; Parkhurst et al., 2013). Additionally, these tissue-resident macrophages possess a high degree of heterogeneity depending on a variety of factors such as the CNS region (van Weering et al., 2023), sex of the organism, stage of life, hormonal exposure, and injury or disease state. Additionally, morphological features can differ depending on microglial interactions with different cellular and non-cellular structures such as neuronal cells, myelinated or non-myelinated axons, and blood vessels in the microglial surroundings and with pathology-associated structures such as amyloid plaques (Baligács et al., 2024; Fujita & Yamashita, 2021; Hattori, 2022).

The dynamic adjustment of microglial morphological features plays a critical role in determining their physiological and pathological functions (Šimončičová et al., 2022). Microglia are equipped with a large number of receptors, enabling them to detect chemical signals, inducing a molecular cascade leading to the modulation of structural properties (McKee et al., 2023). In the healthy CNS, microglia adopt a highly ramified state implicated in the maintenance of homeostasis, for example, through the support of neurons and synapses and surveillance functions (Casali & Reed-Geaghan, 2021; Ueno et al., 2013). Microglia interact with neurons with the release of factors and enzymes from homeostatic microglia such as, for example, the lysosomal enzyme β-hexosaminidase, facilitating neuronal functioning (Frosch et al., 2025). As a result of these neuroprotective functions, the presence of a ramified state is often associated with positive health outcomes in cases of pathology (Vinet et al., 2012). Highly ramified microglia, in the healthy CNS, use their extensive processes and surface-level receptors to monitor their immediate surroundings, scanning for the occurrence of factors posing a threat to CNS health, such as pathogens or signs of injury (Casali & Reed-Geaghan, 2021; Nimmerjahn et al., 2005). When detecting such a threat, such as pathogen-associated molecular patterns located on viruses and bacteria, microglia rapidly retract their processes, which can lead to an amoeboid-like appearance (Casali & Reed-Geaghan, 2021; Nimmerjahn et al., 2005).

Following the encounter with a CNS threat, microglia retract their processes, adopting an amoeboid-like state, associated with the release of proinflammatory cytokines, such as tumor necrosis factor α (TNF-α) or interleukin β (IL-1β), and increased phagocytic activity (Deng et al., 2008). The amoeboid-like morphological state is generally associated with increased motility, enabling the rapid movement towards the site of injury or CNS threat (Lively & Schlichter, 2013). During CNS development, the high level of mobility associated with the amoeboid state enables the dynamic movement of microglia through the CNS and facilitates the pruning of synapses in response to sensory stimulation and the engulfment of apoptotic neurons (Hattori, 2022; Peri & Nüsslein-Volhard, 2008; Tremblay et al., 2010). The increased capacity for phagocytic activity is associated, for example, with the engulfment of degrading cells and cellular debris (Nayak et al., 2014).

In addition to the homeostatic and amoeboid-like states, a large variety of morphological subtypes falling in between these two categories have been identified, undermining the large degree of complexity and heterogeneity of microglial morphological states (Green & Rowe, 2024; Paolicelli et al., 2022). Additionally, in some contexts, in both pathological and health-associated states, a microglial morphological state characterized by a hyper-ramified morphological structure has been identified (Green & Rowe, 2024). This hyper-ramified microglial state may represent an intermediate stage between amoeboid and homeostatic microglia that may precede the adoption of a pathology-associated state (King et al., 2025; Reddaway et al., 2023)

The availability of analysis tools effectively capturing even subtle differences in microglial morphological features across conditions is needed to provide deeper insight into their structure-function relationships across contexts of health, injury, pathology and in response to treatment modalities. Considering the exceptionally high level of diversity and heterogeneity in the microglial population, as well as the dynamic adjustment of microglial morphological features with exposure to a variety of factors, efficient tools to capture this high degree of diversity in terms of morphological features are critical (Tynan et al., 2010). The manual classification of microglia into distinct states can be highly biased, with researchers potentially being guided by pre-existing knowledge regarding differences. An unbiased strategy for the assessment of the presence of morphological states, in contrast, is critical to ensure a high degree of comparability across studies and data sets.

Here we present an easily applicable workflow for the conduction of a clustering analysis to assess the presence and abundance of morphological states in a sample containing microglia, generating distinct cluster assignments for each microglial cell. Subsequently, the abundance of these states in relation to the treatment conditions present in the sample is assessed. The analysis is intended to be conducted following the automated detection of microglia, or other cell types, using a CellProfiler pipeline that was previously developed in the Tremblay lab (Bobotis et al., 2026). The workflow consists of three custom modules to be used within the open-source software CellProfiler. Applying the modules, an outlier assessment, a principal component analysis (PCA) and hierarchical and k-means clustering are automatically conducted, requiring very little user input. The script for the analysis was written in the programming language Python and integrated into modules to be used within the CellProfiler software. The user’s task is thereby limited to the manual adjustment of distinct, sample-specific values. Therefore, the conduction of the proposed clustering analysis requires minimal user input and no coding experience, minimizing the time that will need to be spent on conducting the analysis, while user-friendliness is maximized.

Microglia are characterized by an exceptionally high degree of complexity (Paolicelli et al., 2022). Various measurements can be performed to capture the geometrical and mathematical properties characterizing microglial morphological features. For example, the convex hull describes the area that encompasses an object when a line is drawn connecting the most protruding edges of the object (Graham, 1972). A measurement of an object’s solidity can be obtained by dividing the overall area of the object by the area of the object’s convex hull (Tadic et al., 2019). Compactness is assessed by determining how close the boundaries of the object are located to the object’s center point (Bribiesca, 2008). The bounding box describes the smallest rectangle encompassing the object completely (Rahman et al., 2019). Each of these geometrical assessments provides distinct, valuable information on microglia morphology. By taking into consideration the assessments on a large number of parameters, the high degree of diversity in microglia morphological characteristics can be captured more effectively.

The use of clustering approaches allows for the assessment of the presence and abundance of microglia morphological states, in relation to treatment conditions, by taking into account a large variety of mathematical and geometrical assessments. Through the use of clustering approaches, clusters can be generated, encompassing a subset of microglia characterized by similar morphological features (Kim et al., 2024). Clusters are thereby generated based on the scores of each microglia on a large number of parameters reflecting morphological properties (Kim et al., 2024).

Additionally, by assessing mathematical differences reflecting differences across a high number of geometric properties, morphological subtypes can be assigned in an unbiased manner. This unbiased clustering approach prevents the subjective assignment of microglia to distinct morphological states based on the researcher’s background knowledge.

The conduction of a PCA followed by the use of a clustering algorithm represents a commonly employed workflow for the analysis of patterns representing systematic cellular differences within a sample (Fernández-Arjona et al., 2017). The goal of PCA is to reduce multicollinearity of the included parameters, producing uncorrelated parameters that are linear combinations of the original variables, capturing a large proportion of the variance present in the original dataset (Greenacre et al., 2022). The use of the k-means clustering algorithm enables the detection of distinct clusters characterized by a higher degree of similarity of the data points within compared to the data points between those clusters (Fernández-Arjona et al., 2017).

The necessity to rely on programming languages for conducting clustering analyses limits the wide accessibility of those methods, as at least basic knowledge concerning the functioning of the code is required. Additionally, the necessity to repeatedly adjust a script can increase the risk of errors, potentially slowing down the analysis process.

Therefore, we introduce a user-friendly set of CellProfiler modules that were designed to enable the performance of PCA analysis, followed by a hierarchical and k-means clustering analysis. Additionally, an outlier analysis was integrated that allows the removal of detected microglia that strongly diverged from the remaining detected microglia in terms of their values on the included parameters. To visualize the data distribution and analysis output, interactive plots displaying the cluster assignments and hierarchical structure of cluster assignments within the sample data are generated. As a final output, a bar plot is generated, reflecting the frequencies of the microglia falling into each cluster.

## 2. Methods

### 2.1. Animal and Image Acquisition

This analysis utilized samples from 2-month-old C57BL/6J male mice (Hadera et al., 2025). After purchase, the mice were kept at the Central Animal Facility at the University of São Paulo, Ribeirão Preto campus (Hadera et al., 2025). Animal housing conditions and all procedures were approved by the Ethics Committee of the Ribeirão Preto Medical School (protocol #49/2021) (Hadera et al., 2025). All procedures were performed according to Brazilian and international guidelines on the care and use of research animals (Hadera et al., 2025). Transcardiac perfusions were performed using 4% paraformaldehyde (PFA) in 0.1 M phosphate-buffered saline (PBS). Comparisons were performed between two brain regions: the ventral hippocampus (vHip) and medial prefrontal cortex (mPFC). Sections were selected in accordance with the stereotaxic atlas of Paxinos and Franklin (*Paxinos and Franklin’s the Mouse Brain in Stereotaxic Coordinates*, 2012). The vHip and mPFC are frequently affected across pathologies affecting the CNS, with disruptions in functioning being linked to cognitive and emotional alterations (Belleau et al., 2019; Jobson et al., 2021). The anti-Iba1 primary antibody was used as a marker for microglia (1:1000, Wako, #019-19741) (Hadera et al., 2025). Images were obtained with a Leica LAS X SP8 confocal microscope (LEICA, Wetzlar, Germany) using a 40x objective (numerical aperture=1.4) (Hadera et al., 2025). For each image, z-stacks were acquired, with 19 µm between each plane, generating approximately 20 stacks per image (Hadera et al., 2025).

### 2.2. Image analysis

All acquired images were analyzed using a pipeline created with the CellProfiler software (Broad Institute; v.4.2.6), which can be downloaded here (https://github.com/tremblaylab4-001/CellProfiler-Microglia-Density-Morphology).

CellProfiler modules were added to form a pipeline enabling the automated detection of morphological features (Fig. 1). Additionally, the pipeline facilitates the assessment of these features based on their scoring on a number of parameters capturing geometrical and mathematical differences in the detected objects. Briefly, settings, including a thresholding strategy and value, were adjusted to enable the detection of microglia.

**Fig. 1.**
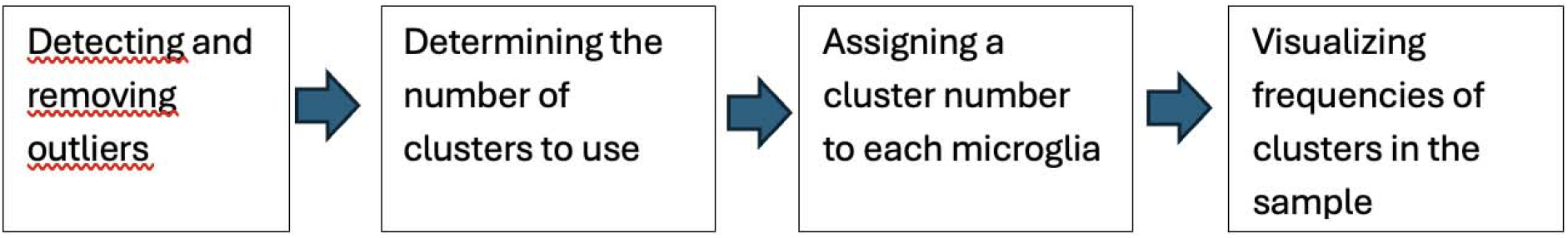
Workflow of data analysis modules. The workflow introduced in this study includes the detection and, following inspection, removal of the outliers. The ideal number of clusters will be chosen, and a cluster number will be assigned to each microglia. Finally, the frequencies of the clusters present in the sample will be displayed in scatterplots and bar charts.

Once optimized, settings were kept consistent across images (Bobotis et al., 2026).

## 3. Steps to perform before conducting the analysis

### 3.1. General information before starting

For the generation of the data used for the clustering analysis that is proposed in this paper, a CellProfiler pipeline for the semi-automated detection of microglia was used. The output generated by this pipeline is used as input for the clustering analysis presented here. As part of the output, a dataset containing measurements of morphological features is generated. The microglial features are assessed using the “MeasureObjectSizeShape” module. The “ExportToSpreadsheet” module in the CellProfiler pipeline generates a spreadsheet containing all the microglial measurements conducted by the pipeline. The name that will be assigned to the spreadsheet can be set in this module. In the spreadsheet, each column represents one parameter reflecting one microglial morphological feature, and each row represents each of the assessed microglia. Additionally, this spreadsheet contains information on the filename of the image in which the microglia have been detected, as well as the number of the microglia (Fig. 2). Within each image, the detected microglia are numbered. Therefore, to identify a specific microglia using the screenshots generated by the CellProfiler pipeline, both the image and object number as presented on the spreadsheet are required. Secondly, the CellProfiler pipeline generates folders containing screenshots of the microglia as identified by the CellProfiler pipeline. Each screenshot contains the coloured shape of the identified microglia on a black background (Fig. 3). These images can be used to compare the location of the microglia in the plots generated as part of the output of this analysis and in relation to other identified microglia. Prior to starting the analysis, a **Condition** column should be manually added to the data frame. This column should contain information on the level of the condition which will be used to conduct comparisons between groups. In the analysis here, comparisons will be made between the two brain regions, therefore, for each image the level of the condition, here, either “vhip” or “mpfc” will be added for each microglia. Additionally, for each image, the blinded number should be present in a column titled **FileName_Blind**. The column containing the number of the specific microglia for each image should be labeled **ObjectNumber**. It is important to assign the exact labels as described here to the columns, as this is the column names will be detected by the code.

**Fig. 2.**
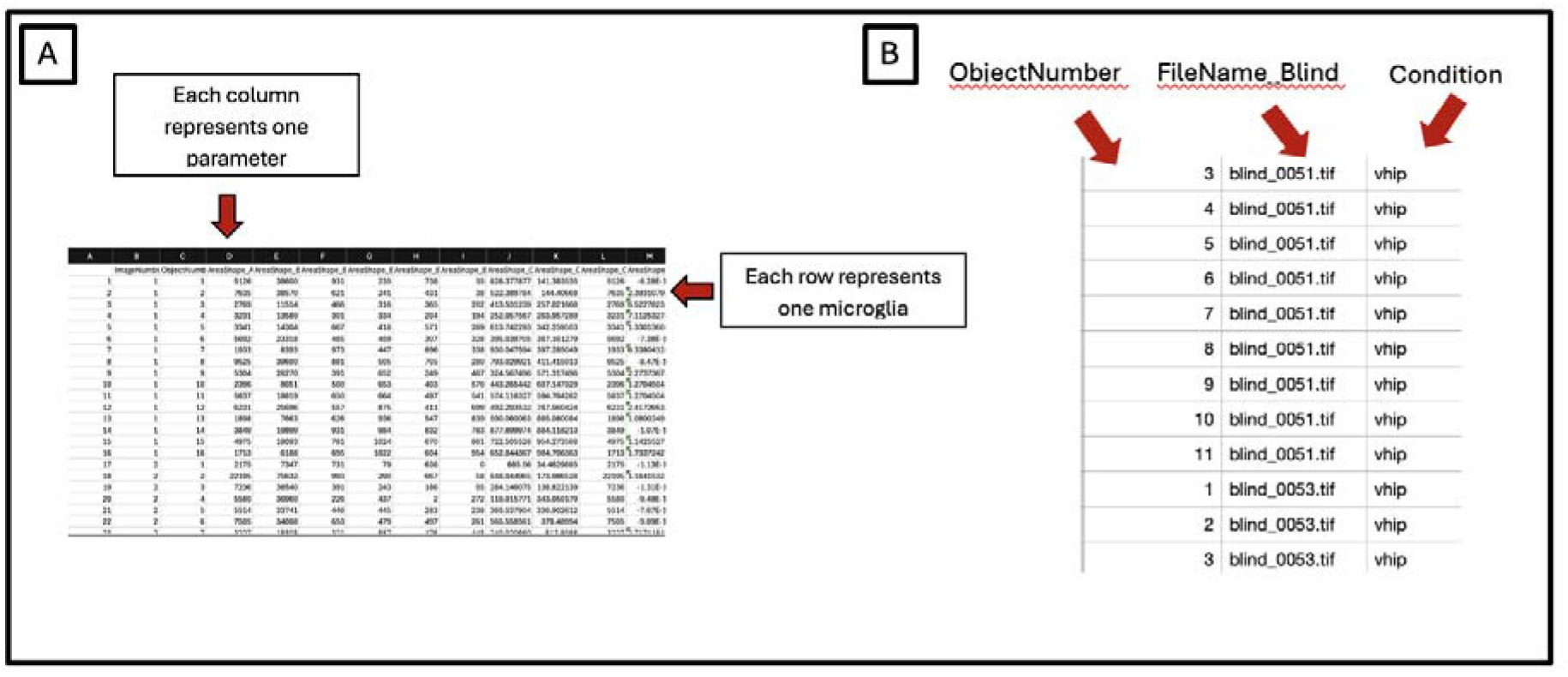
The Input Dataframe. This data sheet constitutes the output data sheet generated by the pipeline containing information on the identified microglia and their corresponding measurements. (A) The data frame contains measurements of microglia morphological features with each column representing one parameter, and each row representing one microglia. (B) A column with the title **Condition** should be added to the data frame prior to starting the analysis, indicating the correct level of the categorical variable for each microglia, here either “vhip” or “mpfc”. Additionally, a “FIleName_Blind” and “ObjectNumber” column should be present.

**Fig. 3.**
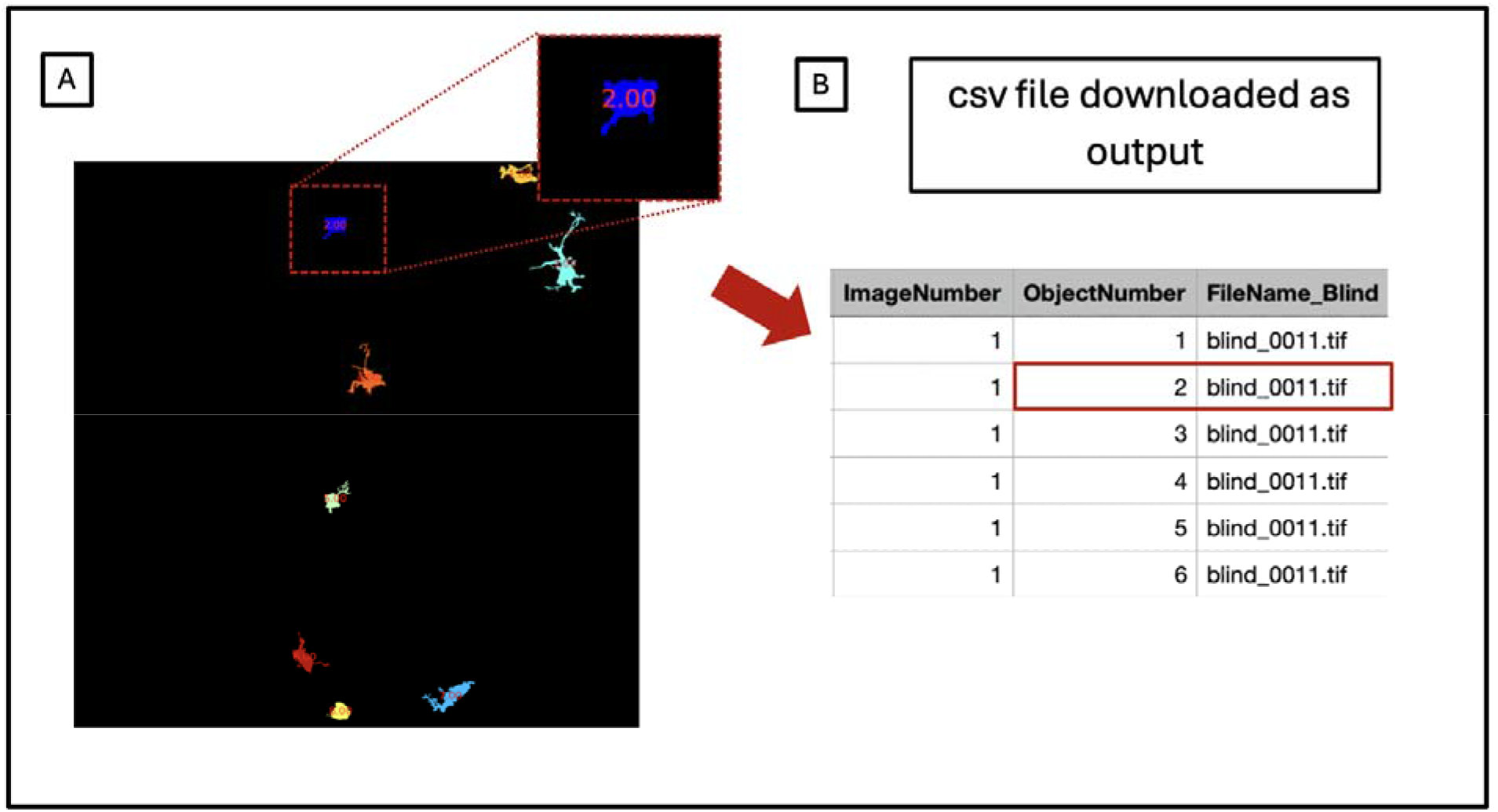
Screenshots showing outlines of the detected microglia. The CellProfiler pipeline generates a spreadsheet as output containing microglial morphological measurements. (A) Images of the objects detected by the CellProfiler pipeline are generated as output by microglial morphology pipelines (Bobotis et al., 2026). (B) As part of the output generated by the custom modules presented in this paper, a spreadsheet contains information on the corresponding filename of the image being generated. The information provided in the spreadsheet can be used to link the object’s position in the scatterplot to the specific object as identified by the microglial morphology pipeline.

### 3.2. Creating the pipeline - loading the custom modules into CellProfiler

Before applying the analysis steps to the data, the custom modules that we created will need to be downloaded from the Tremblay lab GitHub. The workflow required for the download of the custom modules will differ depending on the type of computer that is being used. The setup guidelines corresponding to the computer that is being used for the analysis should be chosen. In the following three sections, instructions will be provided on how to install the CellProfiler modules. Further information on the script that was used to generate the modules can be found on our Github page (https://github.com/tremblaylab4-001/CellProfilerCustomModules).

### Windows setup

1. Before following the subsequent instructions, restart the computer. To get started deploying CellProfiler on Windows, click the Start button, type “Terminal” into the search bar, right-click Terminal and select “Run as administrator” (Fig. 4).
2. 2. Next, copy and paste the following command into the terminal window and hit Enter. ~~~
wsl.exe --install Ubuntu-26.04
~~~
3. Within the terminal, the user will be prompted to create a **username** and a **password**. Note that the letters and characters are <u>not visible</u> while typing the username and password. If a prompt appears asking to restart your computer, restart it. Next, open the terminal again and copy and paste the following line. ~~~
wsl.exe -d Ubuntu-26.04
~~~
4. Next, copy and paste the following block of commands into your Terminal. The user will be asked for the password that was created in the step above; type in the password. Following this step, CellProfiler will automatically open on your computer. ~~~
cd ∼
sudo apt update
sudo apt install -y git libsm6 libnotify4 libsdl2-dev
sudo apt install -y xdg-utils build-essential
curl -fsSL https://pixi.sh/install.sh | sh
. .bashrc
git clone --recursive https://github.com/gnodar01/cp-42x-dev.git
cd cp-42x-dev
pixi add -e dev pandas numpy scikit-learn plotly matplotlib seaborn
git clone https://github.com/tremblaylab4-001/CellProfilerCustomModules.git
mkdir -p ∼/.python3/plugins
cp CellProfilerCustomModules/*.py ∼/.python3/plugins
pixi run -e dev cp
~~~
5. Within CellProfiler, under “File”, choose “Preferences” (Fig. 5A). Under Preferences, the input and output folders, indicating the location of your input Excel sheet and the location where your output documents should be stored, should be indicated. Under Preferences “Default Input Folder” click on “Browse” (Fig. 5B). This step allows you to select the folder where the input csv file will be chosen. After clicking on “Browse”, first select “mnt”, then “c”, then “Users”, your computer username, and within this folder, the file that has your input folder stored (Fig. 5C). Next, choose the same file location that you have selected for “Default Input Folder” for “Default Output Folder”. The documents that serve as the input documents will therefore be stored in the same location as the document downloaded as part of the pipeline.

### Mac setup

To be able to use the CellProfiler module, a line of commands will need to be copied and pasted into the terminal. The terminal can be found on the Mac operating system under “Applications” “Utilities”

1. Open the terminal on your computer.

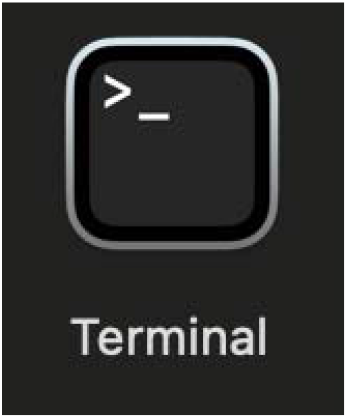

**Fig. 4.**
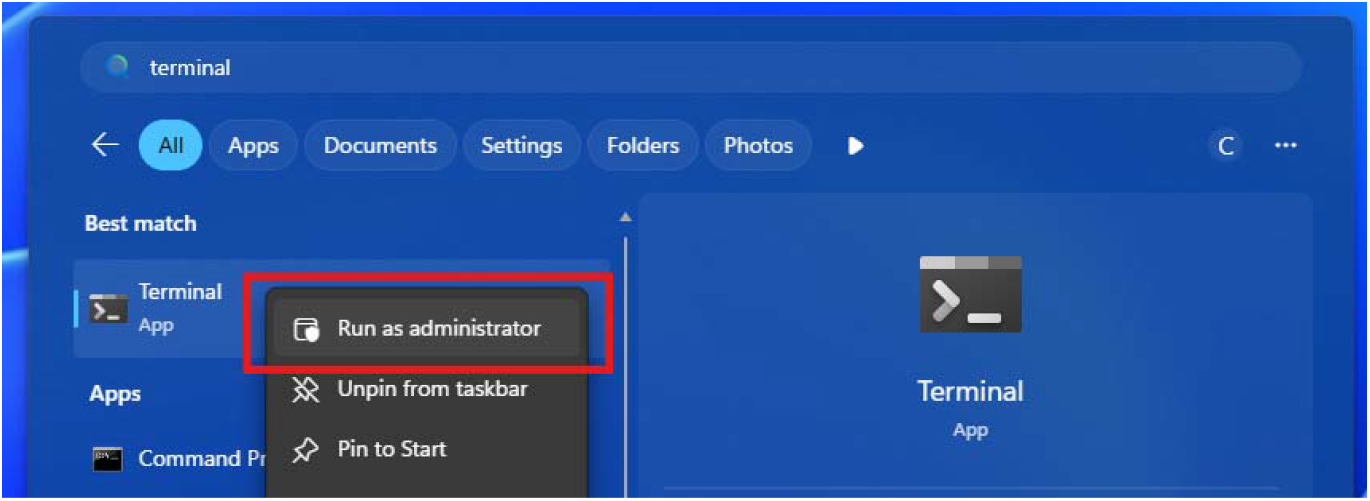
**Terminal location on a Windows computer**. For the setup on a Windows computer, it has to be ensured that the terminal is being run as an administrator.

**Fig. 5.**
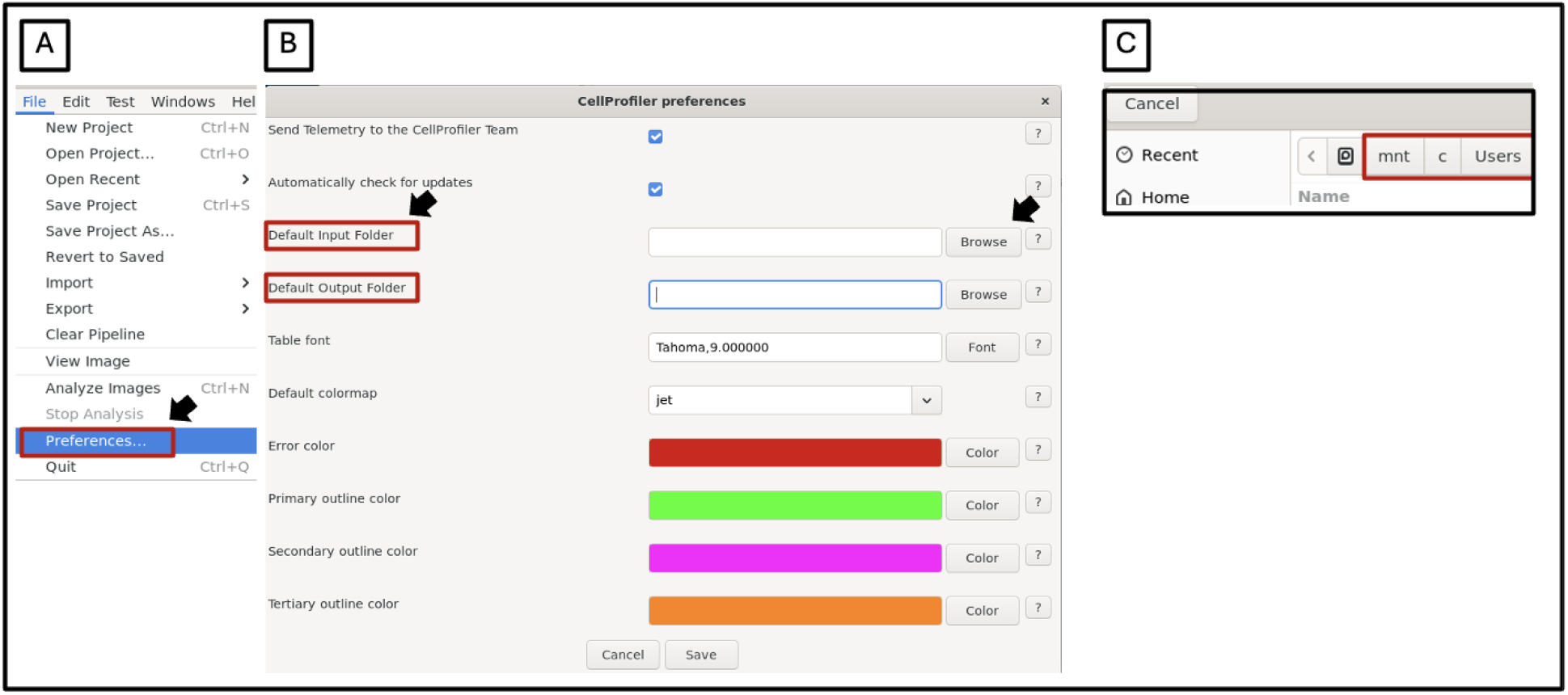
Windows setup - Selecting input and output folders. For users employing a Windows system, within the Preferences window, input and output folders will be chosen. (A) First, within CellProfiler, under “Files”, choose “Preferences. (B) Within the “Preferences” window, input and output folders can be selected. The data frame required for the workflow presented in this paper should be selected as an input. The output folder will determine the folder where the output csv files generated by the modules will be placed following the analysis.

2. Copy and paste the following script into the terminal. Then click “enter” on the keyboard on your computer. Running the code will cause the CellProfiler software to open.

~~~
curl -fsSL https://pixi.sh/install.sh | sh
cd ∼
mkdir cellprofiler-source
cd cellprofiler-source
# creating folder in directory of user xcode-select –install
xcode-select -p
git clone –recursive
# makes folder for cell profiler code
cd cp-42x-dev
pixi install -e dev
pixi add pandas numpy scikit-learn plotly matplotlib seaborn
git clone https://github.com/tremblaylab4-001/CellProfilerCustomModules.git
# download code from github and creating folder containing python #code, putting it
mkdir -p ∼/.python3/plugins
cp CellProfilerCustomModules/*.py ∼/.python3/plugins
pixi run -e dev cp
~~~

1. Within the CellProfiler software, go to “Preferences”. Under “Preferences”, within the “Plugins directory,” delete the directory that appears by default, and copy and paste the following directory information (Fig. 6A). The username can usually be found on your computer under “Users”. At the spot of the “personal username” in the line below, insert the username that is used for your computer. /Users/personalusername/.python3/plugins Under “Default Input Folder” and “Default Output Folder” indicate the folder on your computer used to download the output files into and to pull the input file from. Prior to running the pipeline, your input data frame, containing the morphological measurements in the form of a csv file should be placed into the folder that will be chosen as your “Default Input Folder” (Fig. 6B).
2. After changing the directory information, click on “Save” at the bottom of the page. Then close all CellProfiler windows. To be able to use the modules, the software needs to be closed and restarted first.
3. Next, CellProfiler will need to be relaunched from the terminal; the following command will be needed to be run in the terminal. This command will lead CellProfiler to open again ~~~
pixi run -e dev cp
~~~
4. After closing CellProfiler completely, to be able to use the pipeline again after having already performed the previous steps, every time, before starting, the following command should be entered in the terminal window. This command will enable the user to, once again, have access to the custom CellProfiler modules. ~~~
cd ∼/cellprofiler-source/cp-42x-dev
pixi run -e dev cp
~~~

### Linux (Debian / Ubuntu) setup

## 1. Open a terminal

1. Copy and paste the following commands into the terminal window. The command should lead the CellProfiler software to open on your computer. ~~~
cd ∼
sudo apt update
sudo apt install -y git libsm6 libnotify4 libsdl2-dev
sudo apt install -y xdg-utils build-essential
curl -fsSL https://pixi.sh/install.sh | sh
. .bashrc
git clone --recursive https://github.com/gnodar01/cp-42x-dev.git cd cp-42x-dev
git clone https://github.com/tremblaylab4-001/CellProfilerCustomModules.git
mkdir -p ∼/.python3/plugins
cp CellProfilerCustomModules/*.py ∼/.python3/plugins pixi run -e dev cp
~~~
2. 3. After CellProfiler opens, go to the CellProfiler Preferences. Under the “Plugins Directory,” delete the information that are present here by default, and copy and paste the following information. ~~~
∼/.python3/plugins
~~~
3. Close the CellProfiler software by clicking on the “x”, then re-launch by copying and pasting the following command into the terminal window. The command will lead the CellProfiler software to open again. ~~~
pixi run -e dev cp
~~~

### 3.3. Creating the CellProfiler pipeline

Within CellProfiler, the custom modules can now be added. To add the modules, click on the plus sign next to “Adjust modules”. Modules can be added by clicking on the “+” symbol next to “Adjust modules” (Fig. 7A). The modules to be added can be found under the “Utility” category and should be added in the following order (Fig. 7B).

**Fig. 6.**
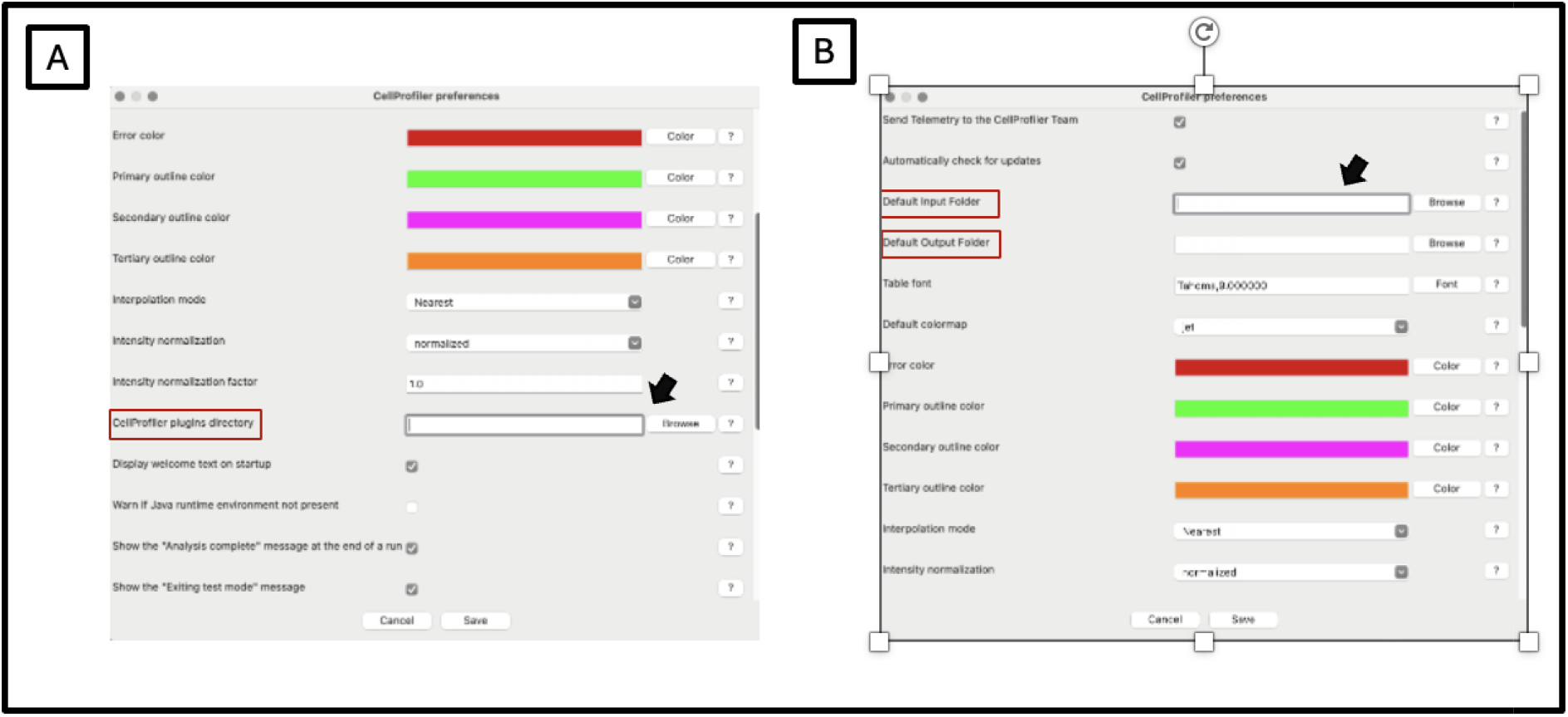
Mac setup – choosing input and output locations. Within the Preferences window, select the location of both your input and output folders. (A) “CellProfiler plugins directors” indicates the location of the script that will be run to perform the analysis steps required for microglial clustering, and should therefore be set to the specific location indicated above. (B) The location of the input and output folder needs to be determined here.

**Fig. 7.**
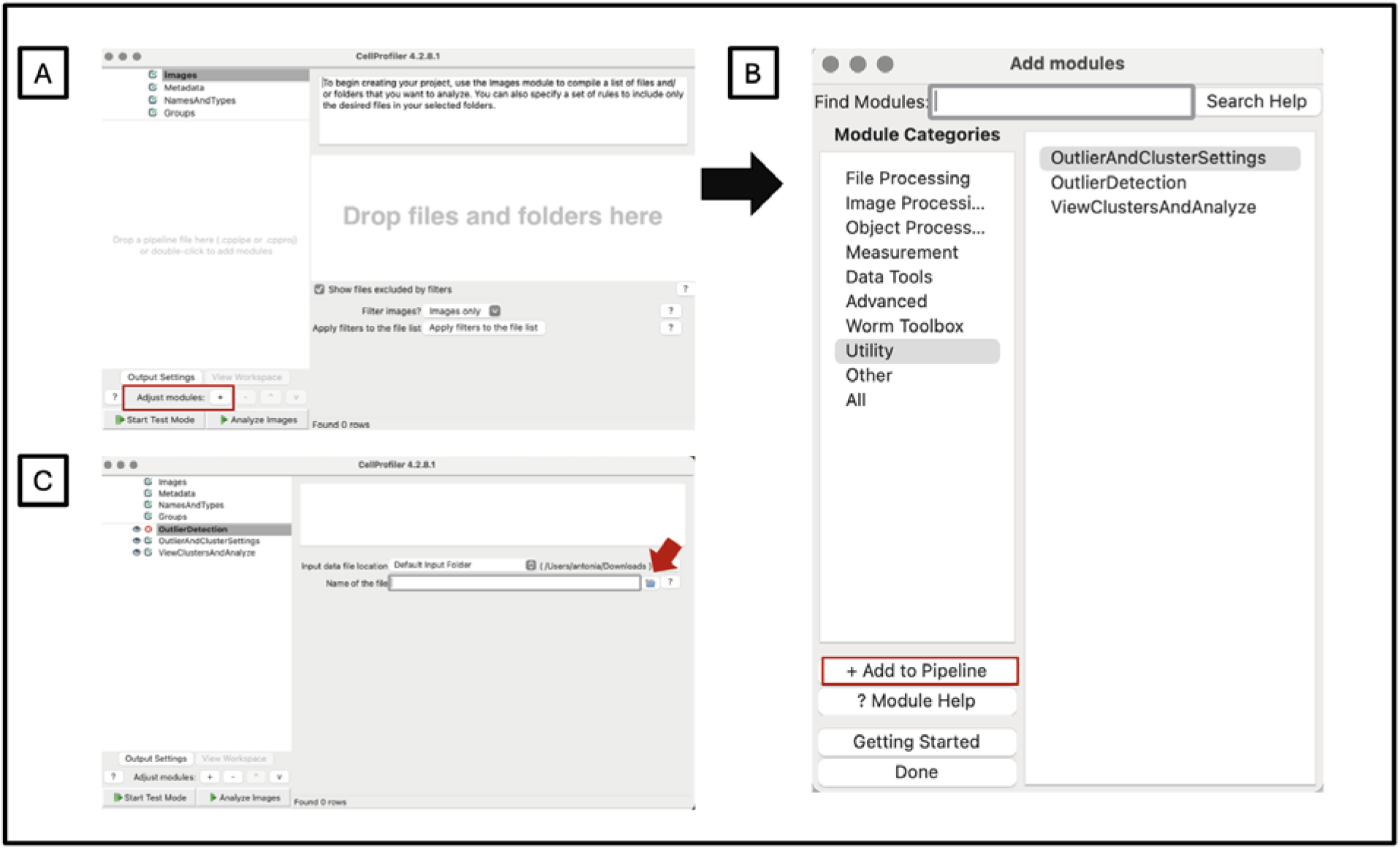
Linux setup – choosing input and output locations. Adding the custom modules to the pipeline. (A) By clicking on the “+” modules can be added to the pipeline (B) The modules used in this presented workflow can be found under “Utility”. (C) The location of the input data frame as determined above, should be selected in this window

“OutlierDetection”

“OutlierAndClusterSettings”

“ViewClustersAndAnalyze”

The three custom modules presented in the pipeline can be run in isolation without any other modules forming part of the pipeline. <u>However, as CellProfiler is a software</u> <u>primarily designed for the analysis of images, to be able to run any pipeline, CellProfiler</u> <u>requires the presence of an image file in the “Drop files and folders here” box within the</u> <u>“Images” module</u>. Before starting the running of the pipeline, it therefore should be ensured that at least one image has been dragged into this box prior to starting. The presence of an image file in this box should be ensured even if the only modules that will be used are the data analysis modules presented in this paper, and no image analyses will be conducted as part of the pipeline. Without any images placed here, the CellProfiler modules will not be able to run.

### 3.4. Running the CellProfiler pipeline

1. As a first step, the data generated by the CellProfiler pipeline will be loaded into the next module by selecting the input folder in the “OutlierDetection” module. The input data frame should be added by clicking on the folder icon under “Name of the file” in the “OutlierDetection” module (Fig. 8).
2. Next, the created pipeline can be run by first clicking on “Start Test Mode” and then “Step” to run the modules one at a time (Fig. 8).

The next sections will include detailed instructions on the use of the modules.

**Fig. 8.**
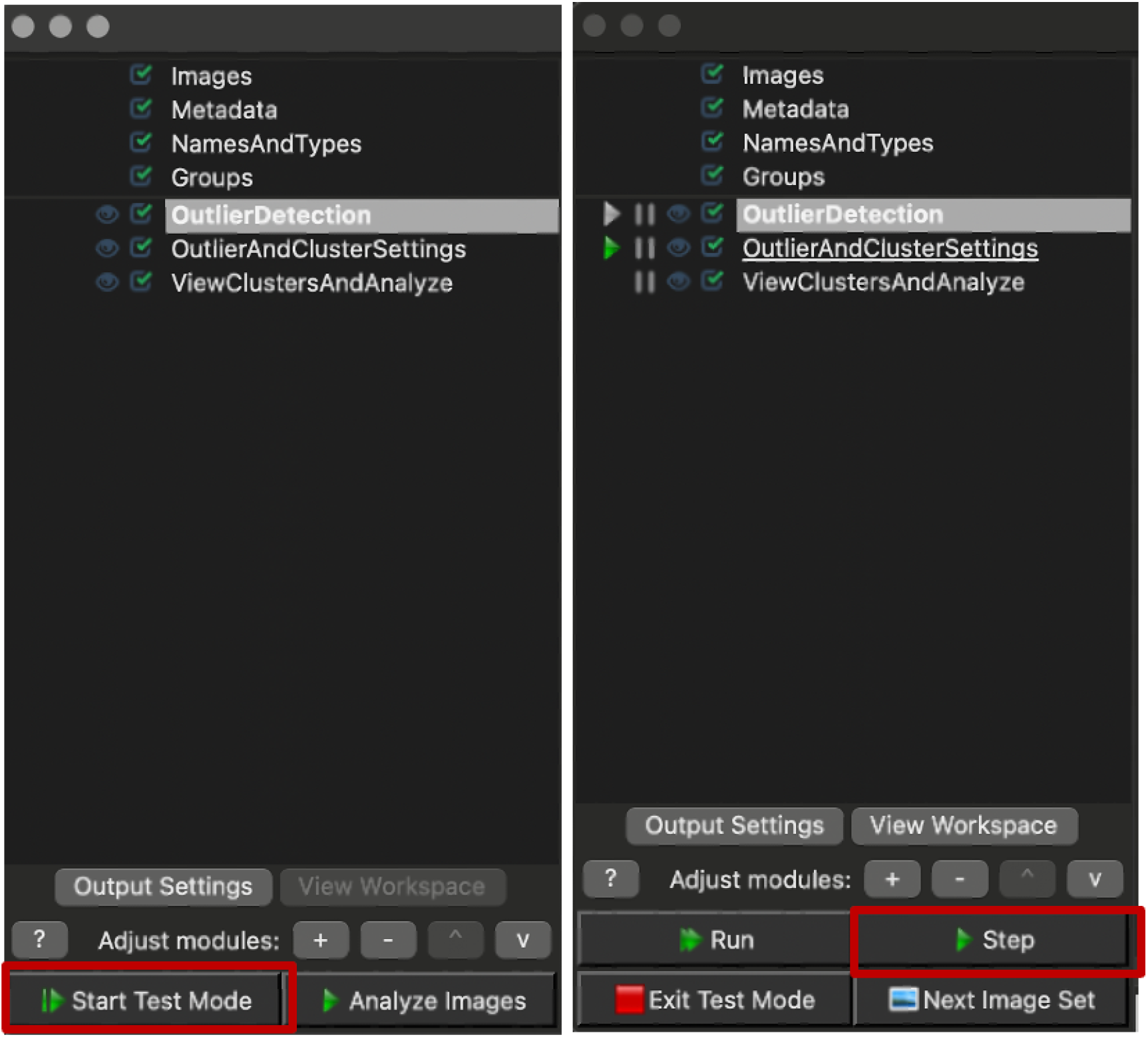
Running the Pipeline. Each module should be run individually using the “step” button.

## 4. Running the Pipeline

### 4.1. “OutlierDetection” module – Checking for outliers

The aim of the outlier detection module is to flag objects in the dataset as outliers that represent artifacts that have falsely been identified as microglia while running the CellProfiler pipeline. The outlying values represent a selection of identified objects that fall outside the overall distribution of data points and therefore deviate strongly from the morphological features assessed in the remaining objects (Alghushairy et al., 2021). The threshold value that is used for determining outlying values has been optimized across several data sets and will not need to be changed between samples.

1. As an output of this module, the user is presented with an interactive three-**dimensional plot**. Each point in the scatter plot represents an object that has been identified as a microglia by the CellProfiler pipeline. The outlying values are coloured in red, and the remaining data points are coloured in black (Fig. 9A). Hovering with the cursor will lead to the appearance of the corresponding label for the specific object, providing information on the image and object number (Fig. 9C). This information can be used to check a screenshot containing the shape of the corresponding object as detected by the CellProfiler pipeline.

2. To check the microglia screenshots corresponding to the objects that have been flagged as outliers, as a second output of this module, **a CSV file is being generated** (Fig. 9B). The CSV file, titled “outliers_dataframe.csv” will be automatically downloaded onto the user’s computer into the folder that has been selected as “output folder” under “preferences” in the set-up. It should be noted, that running the modules again will lead the previously downloaded dataframe to be overwritten by the most recently created dataframe, as it will be downloaded under the same name. Each row in the spreadsheet represents one microglia and contains information on the corresponding image (third column), object number (second column) and the outlier status (fifth column). To ensure that the objects labeled as outliers by the algorithm represent objects that have falsely been identified as microglia by the software, the user should check a screenshot of each of the identified objects individually. CellProfiler provides modules that can be used to create snapshots for the detected cells, which can be used to check the objects corresponding to the labels in the data sheet. As indicated in the steps above, for the images used here, each image name starts with “blind”. The user should check the screenshots of the objects flagged as outliers by the algorithm to decide whether or not to remove the outlying value. The images containing the objects detected as microglia by the pipeline, and that have been used to assess the morphological characteristics of the detected objects in the presented analysis, can be found on our Github ().

3. As part of this module, a small window titled “Edit Outlier Data” will appear (Fig. 9D). As default values, the indices of all the objects (row numbers, see orange arrow (Fig. 9B) that have been identified as outliers within this module are listed here. Clicking “OK” in this window will lead to the automatic removal of all the listed rows from the data set. Once the rows are removed from the data set, the updated data set will be used as input for the next module.

**Fig. 9.**
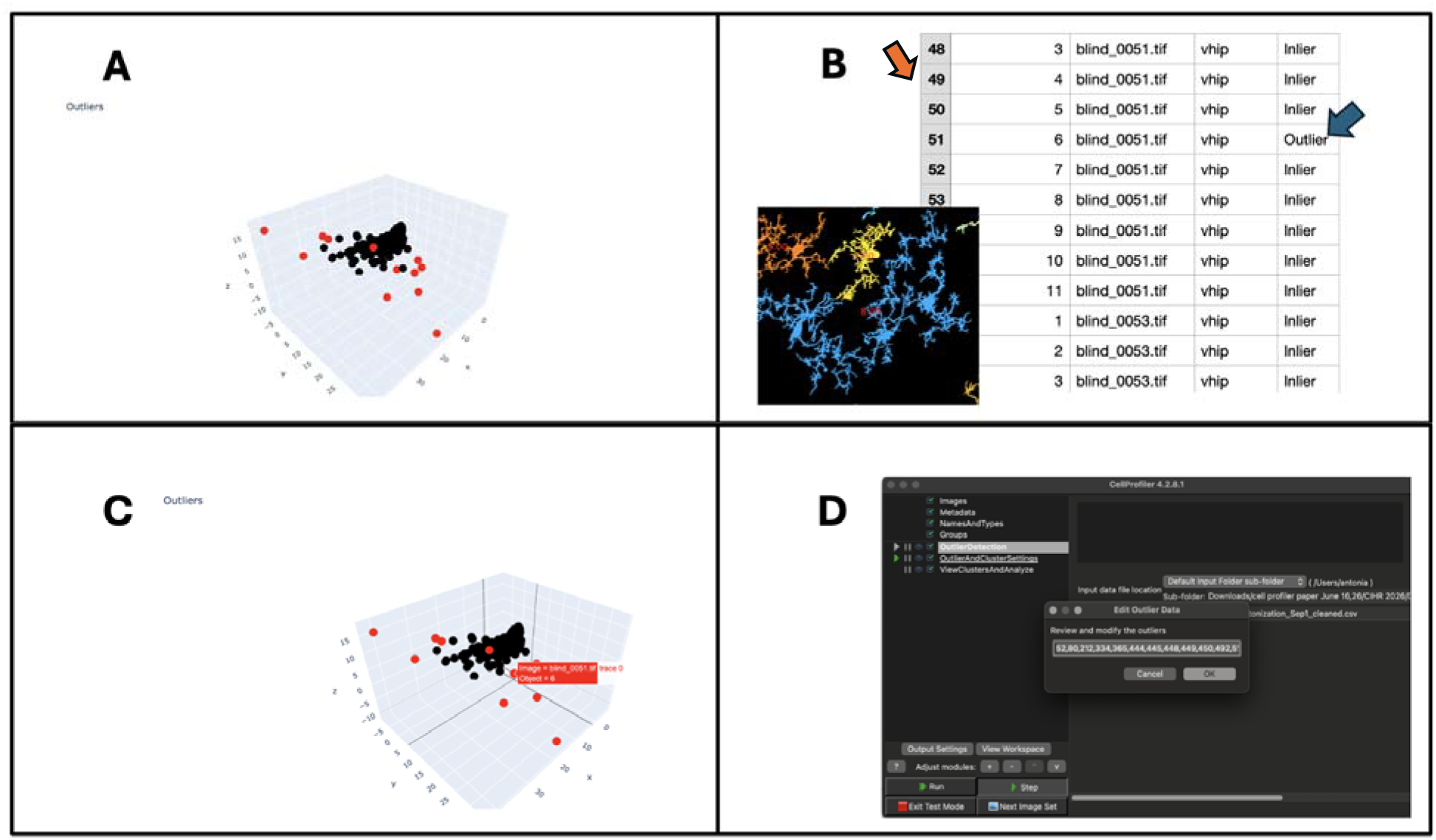
**Outlier module – outputs**. (A) A three-dimensional graph is created; outliers appear in red, inliers in black. (B) A data sheet is created and downloaded onto the user’s computer containing all the identified objects and the corresponding image and object number and outlier status. Each of the labeled outliers should be checked separately by the user to assess whether it represents an artifact and not a microglia. (C) The user can hover the cursor over each of the data points in the graph to receive information on the image and object number. (D) After the user has decided which objects to remove, the values of the index numbers corresponding to the objects representing artifacts to be removed can be placed in the “Edit Outlier Data” window. The index numbers for the corresponding cells can be found in the Excel sheet displayed in panel 2 (indicated by the orange arrow). By default, a list of all the outlying values appears in the box, and the user will have to manually remove the index of the object/ microglia that should be kept as part of the analysis.

### 4.2. “OutlierAndClusterSettings” module – Choosing the number of clusters in the sample

To determine the number of clusters that best reflect the differences in microglia morphology present in the sample, a hierarchical clustering analysis is applied. The hierarchical clustering reveals the data structure reflecting the groupings based on morphological similarities present in the data (Murtagh & Legendre, 2014). As an output of the analysis, a dendrogram is created, revealing a hierarchical display of the clusters at different levels of similarity (Nielsen, 2016). The dendrogram displays groupings starting from the level of individual microglia at the bottom, repeatedly grouping similar individual clusters together, until only one cluster remains at the very top (Nielsen, 2016). Including a larger number of clusters enables a better capture of the unique morphological features present in the identified microglia, reflected by a smaller within-cluster sum of squared errors. In contrast, selecting a smaller number of clusters leads to more prominent morphological characteristics shared by the instances in the cluster driving the clustering assignment, but is, however, associated with a higher error rate.

1. This module generates a dendrogram as an output that will inform the user of the microglia clusters present in the sample and help determine the ideal number of clusters to include (Fig. 10A).
2. Following the generation of the dendrogram, the user can decide on a threshold value. A small window appears as part of the module prompting the user to type in a threshold value. We suggest that a threshold around the 50^th^ to 60^th^ percentile of the highest Ward’s distance (highest Y-axis value) present in the sample should be selected. Additionally, the same threshold should be applied across all analyses to facilitate an increased level of comparability across samples. The user input will determine a threshold that will be presented as a line on the dendrogram. The threshold will represent the location where the dendrogram is being cut determining the number of clusters that will be created. Based on the threshold that is being used, the immediate groups below that threshold will be displayed in different colours (Fig. 10B). Further information on hierarchical clustering and the generation of the dendrogram can be found under supplementary materials.

**Fig. 10.**
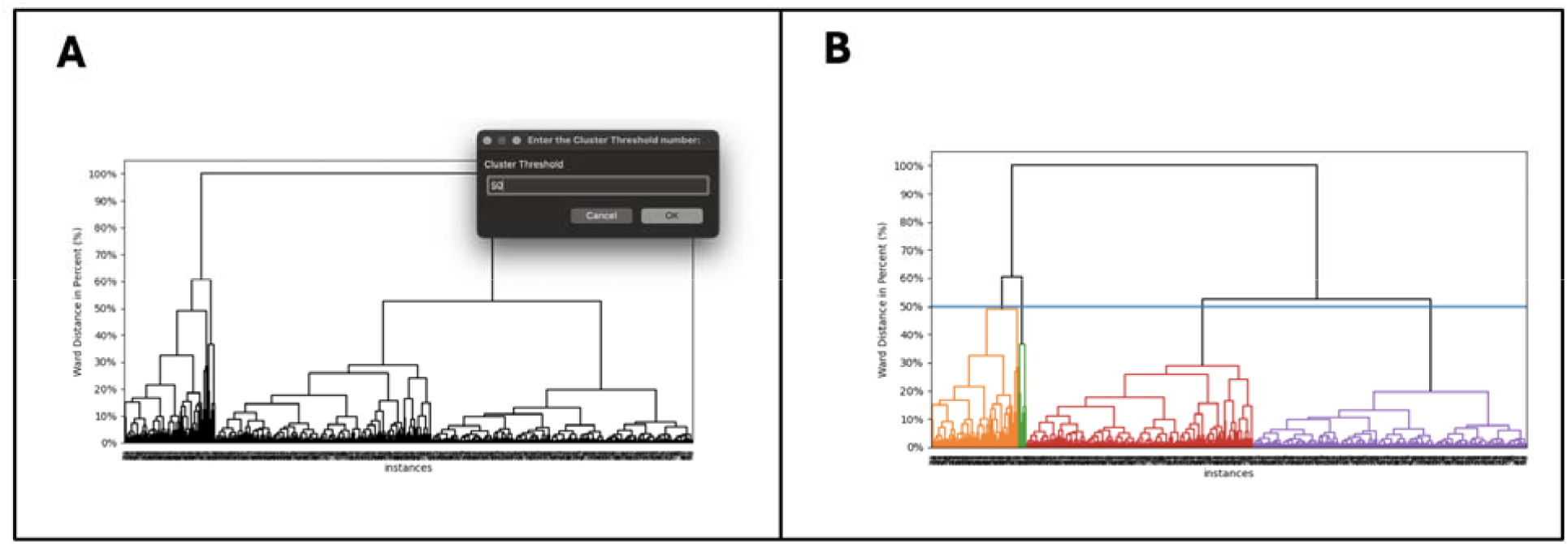
Choosing the number of clusters in the sample. The user receives an image displaying a dendrogram as output. The individual lines at the very bottom level of the plot represent individual microglia. The clusters are repeatedly merged until only one cluster remains at the very top of the plot. The clusters at different heights represent groupings at different levels of similarity, with the central point in each cluster differing more strongly at higher levels. (A) Based on the graph, the user can decide on a threshold (reflecting the Ward distance value, in percentage, on the Y-axis). The dendrogram can be used to guide the decision on the number of clusters to choose. (B) The created clusters are presented in distinct colours in the modified dendrogram. Based on the user input, the immediate clusters present below that threshold will be visualized using a different colour (Fig. 10,B). The user will then receive an image containing the same dendrogram, with each of the clusters visualized in a different colour.

### 4.3. “ViewClustersAndAnalyze” module – Assigning each microglia to a cluster

After deciding on the number of clusters to include in the module above, each of the individual microglia will be assigned to a cluster. Additional information on the use of k-means clustering to assign each microglia to a cluster can be found under supplementary materials.

1. Microglia within the same cluster are characterized by a higher degree of similarity in terms of morphological features. A three-dimensional plot is generated visualizing the distribution of the individual microglia in terms of morphological differences. Each microglia is displayed in a colour representing its cluster assignment.
2. As part of this module, a three-dimensional plot will be generated reflecting the distribution of the clusters (Fig. 11A)
3. The next step is to identify how the clusters differ in terms of the morphological features characterizing each cluster. To examine the morphological features present within each cluster, the next step will be to examine the screenshots of microglia falling in the different clusters. Hovering with the cursor over each data point in the graph will lead to the appearance of a label containing information on the corresponding image and object number (Fig. 11B). Based on this output, the user can trace back the individual microglia (represented by individual rows in the data sheet) to examine the corresponding screenshot (Fig. 11C-D).
4. The user will further receive a data sheet as an output that will contain the microglia condition, image number, object number and cluster assignment (Fig. 11C-D).
5. Following the window displaying the interactive three-dimensional cluster graph, an additional window will appear displaying a bar plot (Fig. 12). The bar plot visualizes the number of microglia in each of the clusters for each level of the condition included in the analysis.

##### Box 1: Interpretation of the clusters – Detection of outliers grouped into the same cluster

The clustering analysis groups microglia resembling each other in terms of morphological features into the same cluster. However, in some cases, the clustering might group together artifacts that have been falsely identified as microglia. For example, one cluster may consist of several objects consisting of multiple microglia that have been identified as a single object. In this case, the clustering analysis provides additional insights into the existence of outlying values. If the outlying values are systematically grouped together into the same cluster, the corresponding rows should be removed from the input data sheet in the first step, and the analysis should be repeated. However, to prevent the formation of clusters representing artifacts, the user should ensure when running the pipeline that the detected object represents a microglia.

**Fig. 11.**
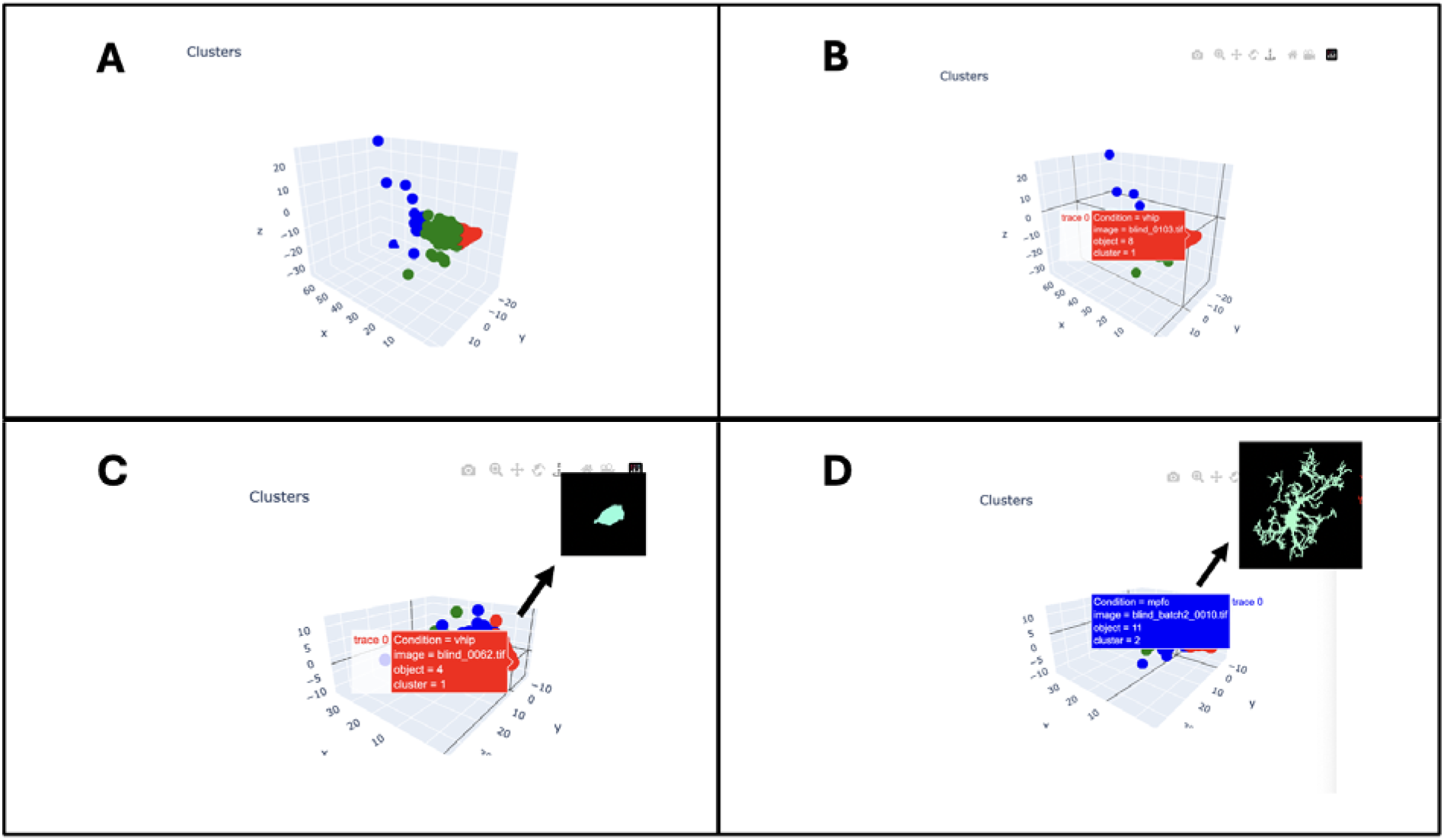
Determining the morphological features characterizing each cluster. After the user has determined the number of clusters in the previous step, each microglia will be displayed in a three-dimensional plot and displayed in a colour representing its cluster assignment. (A) Each of the three identified clusters is represented in a different colour. (B) Hovering over each of the data points in the scatterplot will lead to the appearance of a label containing information on the image and object number. (C) The information displayed on the label can be used to check a screenshot of the identified microglia, each of which can be found on a numbered image, and possesses an individual number within that image. (D) For each of the clusters a number of representative microglia should be checked to gain a deeper understanding of the morphological characteristics of the microglia within a specific cluster.

**Fig. 12.**
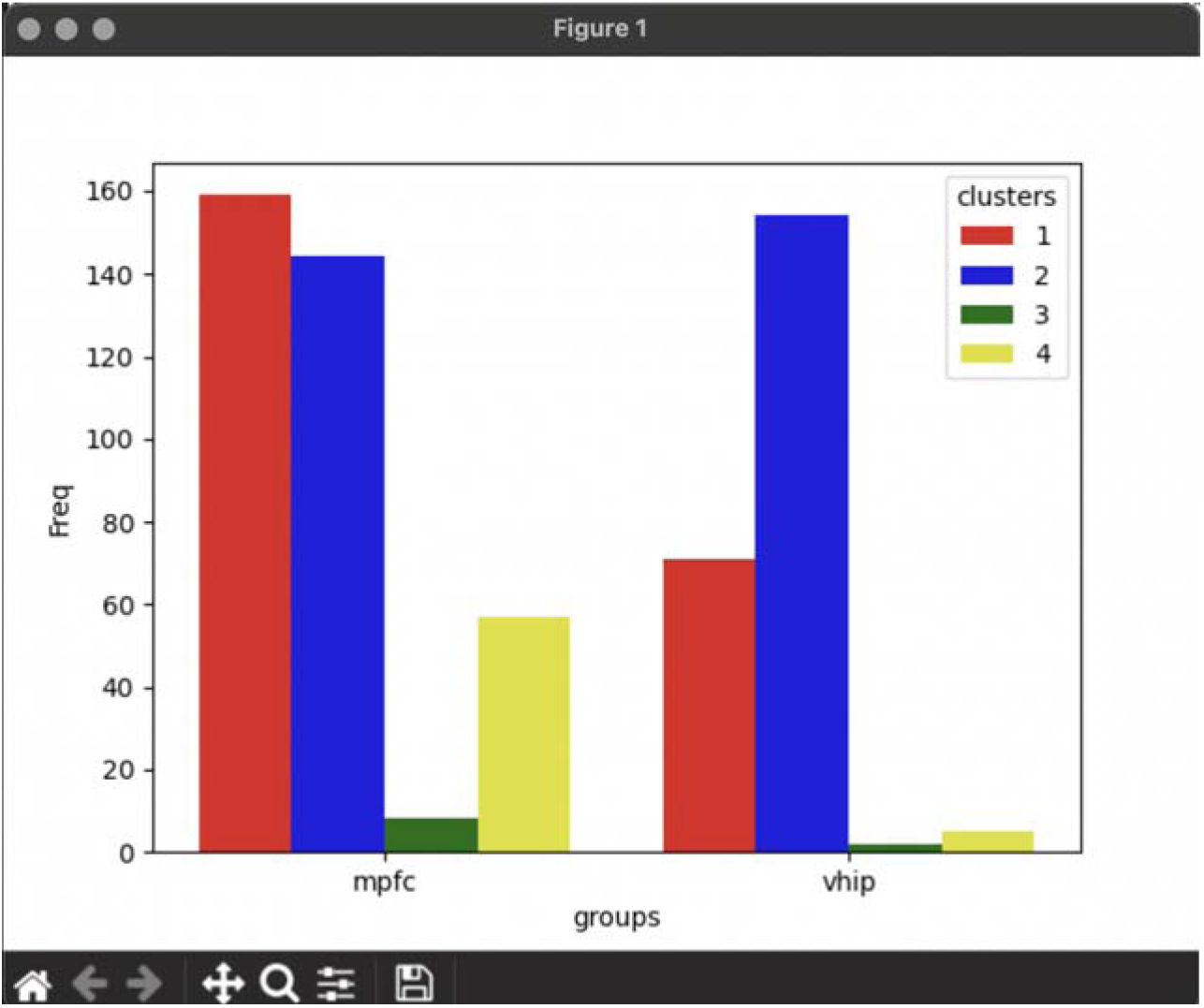
Creation of a bar chart as output. A bar chart will be created displaying the frequencies of the clusters present within the conditions.

**Fig. 13.**
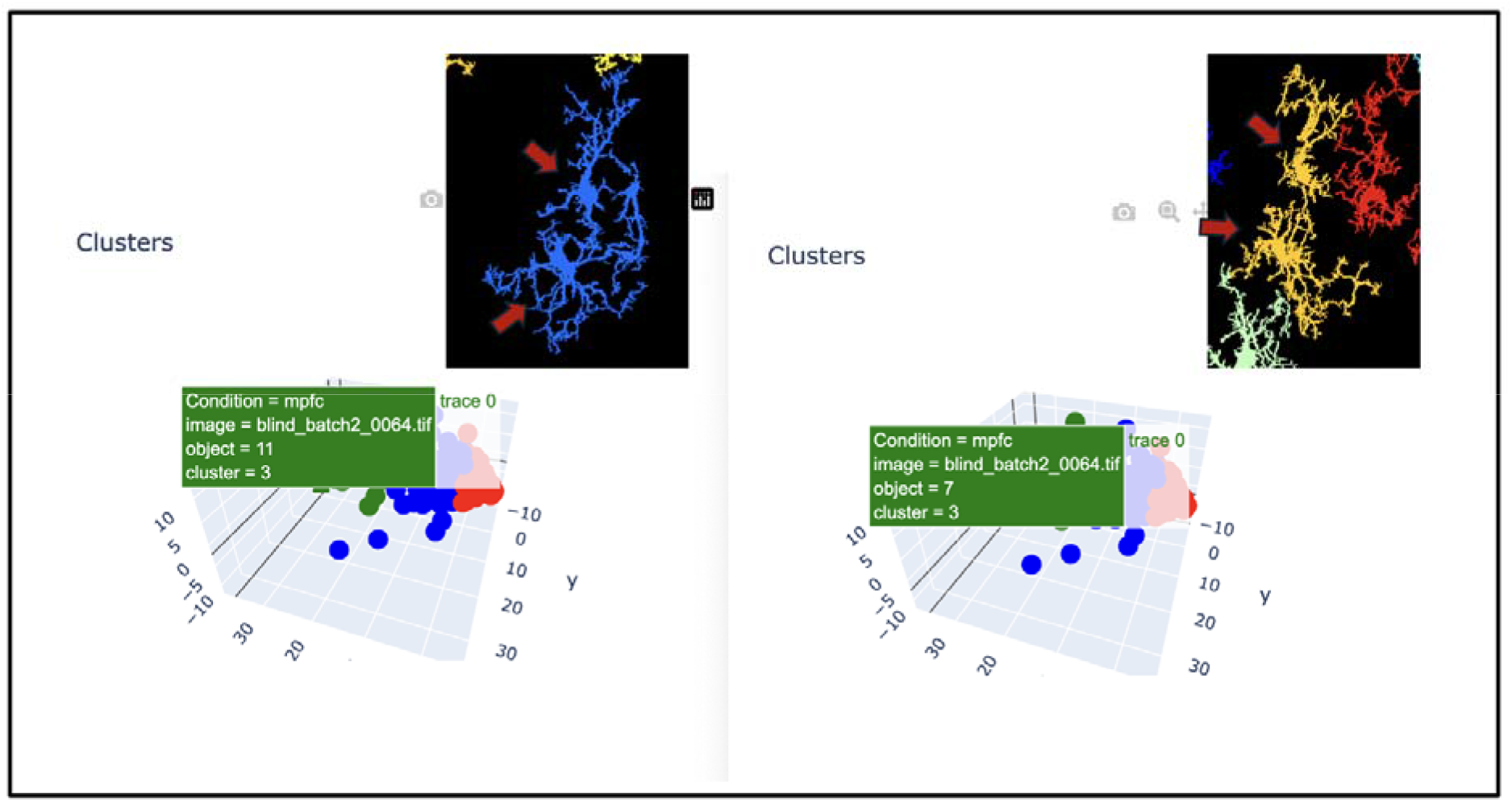
Detection of outliers grouped into the same cluster. Example of a graph displaying a clustering analysis grouping together several clumped objects into the same cluster. Several objects within the same cluster consist of two separate microglia that have been identified as the same object. The red arrows indicate the cell bodies of the separate nuclei. The labels provided in the 3D plot indicate the corresponding row in the input data sheet that should be removed from the dataset.

## 4. Discussion

Microglia are a highly dynamic group of cells, with their structural diversity reflecting the heterogeneous functions they fulfill (Green & Rowe, 2024). Understanding microglial morphological adjustments across states of physiology and pathology, and in response to different stimuli, such as pharmacological and non-pharmacological treatments, will provide valuable insights aiding in the development of efficient therapeutic approaches (Šimončičová et al., 2022). Therefore, to gain a deeper understanding of the role of this cell type in the mechanisms underlying the efficacy of different approaches, the availability of efficient strategies enabling the capture of this high degree of diversity is critical.

Here we present a set of custom modules for the open-source software CellProfiler enabling the clustering of microglia based on differences in their morphological features. The presented CellProfiler modules enable the application of hierarchical and k-means clustering algorithms following a PCA analysis to divide the microglia in the sample into clusters. The clusters are thereby created to minimize the differences within the clusters and maximize the differences across clusters to best capture biologically meaningful differences in cell morphology. Before these analyses, an outlier test can be conducted, enabling the exclusion of detected objects that fall far outside the distribution of the remaining objects.

The workflow that we created in the form of three distinct custom CellProfiler modules, requiring minimal user input, includes the conduction of a PCA, an outlier assessment, and the application of hierarchical and k-means clustering approaches. As the parameters measuring microglia morphological features are often highly correlated, before the conduction of a clustering approach, a PCA was conducted. A PCA enables the linear recombination of a set of included variables, leading to the generation of a set of uncorrelated features capturing the majority of the variance of the originally included variables (Greenacre et al., 2022). Therefore, a PCA leads to the generation of a set of uncorrelated parameters reflecting the scoring of each microglia on the original set of variables. The PCA output can therefore be used as input to the clustering analysis (Fernández-Arjona et al., 2017).

As part of the presented analysis workflow, an easily applicable outlier assessment that will limit the inclusion of artifacts into the clustering analysis is presented. The application of clustering analyses is highly sensitive to outlying values (Gan & Ng, 2017). Especially in automated cell detection pipelines, artifacts, falsely identified as cells, can easily alter and invalidate analysis results. Outliers may represent objects that reflect background staining in immunofluorescent images, objects that consist of multiple cells clumped together into one object, loose processes not connected to a cell body, or only partly visible microglia. The inclusion of outliers into the clusters can distort the analysis, increasing the variance within each cluster (Gan & Ng, 2017). Alternatively, the clustering of a group of similar objects, representing artifacts instead of microglia, such as a cluster consisting of only loose processes, would reflect a cluster not representing a microglial morphological states, leading to the distortion of results.

Therefore, before the conduction of a clustering analysis, an outlier test was integrated as a separate analysis step. The included outlier test includes the calculation of a local outlier factor (LOF) for each of the individual identified microglia (Alghushairy et al., 2021). The integration of this highly user-friendly outlier test prevents the use of a data set as input that is distorted by individual, falsely detected artifacts that may decrease the reliability and validity of the clustering output.

Following the conduction of the outlier assessment, before assigning each microglia to a cluster, a hierarchical clustering is conducted. To be able to assign each microglia to a cluster using a k-means clustering approach, as is being done here, it is necessary to first determine the ideal number of clusters that should be used to best represent the morphological differences present in the sample. Conducting a hierarchical clustering analysis, followed by the generation of a dendrogram, will provide insights into which cluster number should be used ideally for the sample set at hand (*Cluster Analysis in Marketing Research: Review and Suggestions for Application - Girish Punj, David W. Stewart, 1983*, n.d.). Specifically, the diagram will provide insights into how the inclusion of cluster numbers will affect the within-cluster variance if a cluster is added or removed. Based on the diagram, a number of clusters can then be chosen that best represents the data structure present in the sample.

Determining the number of clusters based on the output provided by the hierarchical clustering analysis will provide useful guidance, enabling the user to make an informed decision for determining cluster numbers. A threshold that can be used as a cutoff value to determine the number of clusters is proposed to increase the comparability across samples. Including this more guided approach for choosing the number of clusters to include will increase the reproducibility and comparability across studies. Another frequently used method for determining the number of clusters to include is through the generation of plots, such as scree plots (Streiner, 1998). Using these plots, the number of clusters to include is determined based on the change in the slope of the line displayed in the graph (Streiner, 1998). The point in the graph characterized by the most drastic flattening in the slope is used to determine the number of clusters. However, this approach can be highly subjective and may therefore limit the reproducibility across studies (Streiner, 1998). By conducting a hierarchical clustering analysis and setting a range of values to be used as guidance to determine the number of clusters to include, we provide an objective and reproducible method for determining the number of clusters (Streiner, 1998).

The presented workflow was created to be employed in combination with existing CellProfiler pipelines. Published CellProfiler modules enable the fast and automated detection of different cell types, including microglia, but also other cell types such as interneurons, assessing their morphological characteristics. However, the modules that we present here could be used in combination with analysis strategies using different software. For example, analysis strategies have been presented to facilitate the detection of microglial morphological features. For example, the use of the IMARIS software enables the detection of microglia in 3D, detecting protruding microglial processes across several planes, allowing for reconstruction into a three-dimensional object (Murray et al., 2024). The presented code could be used in combination with other analysis strategies, provided the software used produces a data set as an output possessing the same format as the input data set used here, including a set of parameters that assess cellular features (as columns) and a set of microglia (as rows) measured along these parameters.

The modules that we provide are highly user-friendly, requiring minimal user input and provide interactive graphs as outputs that provide a visual representation of the distinct morphological groups present in the sample. The presented workflow therefore represents a strategy to assess the presence of microglial morphology clusters in a manner that is fast, reproducible, easy to apply, adaptable and highly user-friendly.

Employing this approach will facilitate the detection of clusters representing microglial morphological differences, limiting the time required for the analysis, while enabling a high degree of comparability across studies.

## Acknowledgements

We acknowledge and respect the Lək□□əŋən (Songhees and X □sepsəm/Esquimalt) Peoples on whose territory the university stands, and the Lək□□əŋən and W □SÁNEĆ Peoples whose historical relationships with the land continue to this day. We acknowledge the input and valuable feedback provided by Dr. Haley Vecchiarelli. We are also grateful to Dr. Adriano Jose Maia Chaves-Filho for contributing to the initial discussions about the workflow presented here.

M-ÈT holds a Tier 1 Canada Research Chair in *Neurobiology of Healthy Cognitive Aging* (CRC-2024-00155). This work was supported by a project grant from the Canadian Institutes of Health Research (CIHR (#PJT461831) and a Natural Sciences and Engineering Research Council of Canada (NSERC) Discovery grant (RGPIN-2024-06043) awarded to M-ÈT. AL was supported by a graduate grant from the Branch Out Neurological Foundation. VH and FVG are supported by the São Paulo Research Foundation (25/09388-9 to VH and 24/20348-6 to FVG).

