## Supplementary materials for "A Novel Open-Source CellProfiler Pipeline for Automated, User-Friendly Hierarchical and K-Means Clustering of Microglial Morphology"

A script was written using the programming language Python. Using the CellProfiler wiki page, the script was adjusted, enabling integration into CellProfiler modules and enabling the creation of user interfaces for each analysis step. To use self-written code to create a CellProfiler module, the code has to be structured in a very specific way, and certain criteria have to be met. Specifically, predetermined function names and properties are required to enable the loading of the modules by CellProfiler. More information can be found here (<https://github.com/CellProfiler/CellProfiler/wiki/Module-structure-and-data-storage-retrieval>).

The following sections contain detailed explanations of the analyses performed within each module. The Python script can be found on our GitHub page (https://github.com/tremblaylab4-001/CellProfilerCustomModules).

Data pre-processing

Before applying the PCA and clustering algorithms, the data will be standardized (Gewers et al., 2021). The CellProfiler output data sheet is not standardized, and, therefore, standardization is necessary to prevent a distortion of results as a consequence of the different scales used for the different measurements (Gewers et al., 2021). Standardizing the data will ensure that the included variables will not differ in terms of the weight they contribute to the overall variance (Gewers et al., 2021).

Preliminary PCA

Next, a PCA will be conducted prior to the removal of the outliers, and once again, once the outliers have been removed. The PCA will be conducted to reduce the large number of input parameters to a smaller number of uncorrelated variables reflecting the entirety of the variance of the original dataset (Greenacre et al., 2022). As all the measurements are conducted on the same cells, it is likely that the measurement parameters are highly correlated (Greenacre et al., 2022). By conducting a PCA, this multicollinearity will be removed (Greenacre et al., 2022). The variance that is common to all parameters will be reassigned, and the original variables are reconfigured into a set of linear principal components (Greenacre et al., 2022; Vidal et al., 2016). A set of PCs will be chosen to maximize the explained variance representing the unique contribution of all the variables that were originally included (Greenacre et al., 2022). To decide on the number of PCs to include, the Kaiser criterion will be used (Greenacre et al., 2022).

Assessment of outliers

The local outlier factor (LOF) algorithm provides a technique enabling the detection of data points that deviate strongly from the density of the data points in its local environment (Alghushairy et al., 2021). The algorithm assesses whether a specific data point is considered an outlier when taking into consideration its relation to the data points in its local surroundings (Alghushairy et al., 2021). The threshold used in this script was chosen after optimizing the settings for this algorithm by applying it to several different samples, until the objects on the border of the samples, furthest away from the centroid of all data points, were flagged as outliers.

PCA

Following the removal of the outlying values, once again, a PCA will be conducted. As the PCA output is impacted by the presence of outliers, a preliminary PCA was conducted above and is now repeated following the removal of the outliers. The Kaiser criterion will be applied, and the number of components to keep will be determined (Greenacre et al., 2022).

Hierarchical clustering

The cluster assignment will be generated through the application of a k-means clustering algorithm (Aggarwal, 2014). However, to apply the k-means clustering, the number of clusters to use needs to be determined in advance (Aggarwal, 2014). To best determine the number of clusters to select, a hierarchical clustering will be conducted, reflecting the groupings of microglia present in the data (Nielsen, 2016). Following the conduction of the hierarchical clustering, a dendrogram will be created that reflects the data structure and the grouping of sub-clusters at different levels (Supplementary Fig. 1) (Aggarwal, 2014). The clusters are created using Ward’s linkage distance. Clusters are created to minimize the variance within each cluster, the sum of squared errors (SSE) (Nielsen, 2016). The SSE represents the squared Euclidean distance, the squared difference between two data points (Aggarwal, 2014). At higher levels, characterized by fewer clusters, there is a higher increase in the total within-cluster variance (Nielsen, 2016). At each merging step, the merging of the clusters occurs in a manner minimizing the increase in the total SSE (Nielsen, 2016). The dendrogram reflects the number of clusters in relation to the Ward’s linkage distance (Y- axis), representing the increase in total variance after merging the clusters from the previous step (Nielsen, 2016).

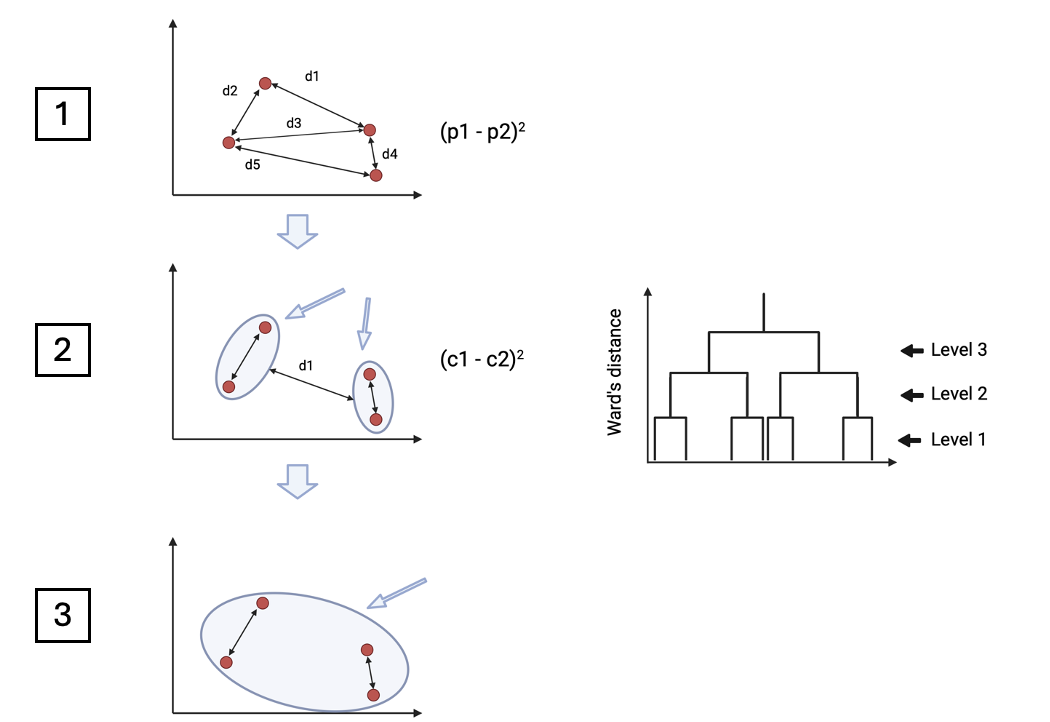

**Supplementary Fig. 1**. **The distance between clusters is calculated iteratively.** First, for each of the data points, representing individual objects identified as microglia, the distance to all the other data points is calculated. Next, each data point is grouped together with its closest other data point, minimizing the sum of squared errors within each cluster. The clusters are repeatedly grouped together as the Ward’s distance value increases until only one group remains at the highest level of Ward’s distance. (1) Euclidean distance values are calculated between each pair of points. (2) As a second step, the two closest points are grouped. The amount of variance that is explained by the fusion of clusters increases with each merge. (3) The clustering is repeated until only one large cluster remains. (4) The dendrogram visualizes the clustering pattern of the data points at different levels of Ward’s distance, reflecting the increase in variance explained between different cluster numbers included.

K-means clustering module

Each microglia is assigned to a separate cluster based on its morphological characteristics. The application of k-means clustering requires the number of clusters to be chosen in advance (Aggarwal, 2014). As a result of the clustering, objects possessing a high degree of similarity, reflected in the numerical distance between the objects, will be grouped, while objects characterized by a higher degree of dissimilarity will be grouped in different clusters (Supplementary Fig. 2) (Sinaga & Yang, 2020). As a first step, for each cluster, an initial centroid location is randomly chosen, and each data point is assigned to the nearest centroid (Supplementary Fig. 2A) (Aggarwal, 2014). Here, the distances are calculated using the Euclidean distance. All difference values, representing the difference from the data points to their centroids, are squared, leading to the generation of within-cluster sum-of-squares values representing the sum of squares explained by all the clusters (Aggarwal, 2014). The goal of the k-means algorithm is to repeatedly recalculate the centroid location to keep within-cluster sum-of-squares values as small as possible (Supplementary Fig. 2B) (Arthur & Vassilvitskii, n.d.).

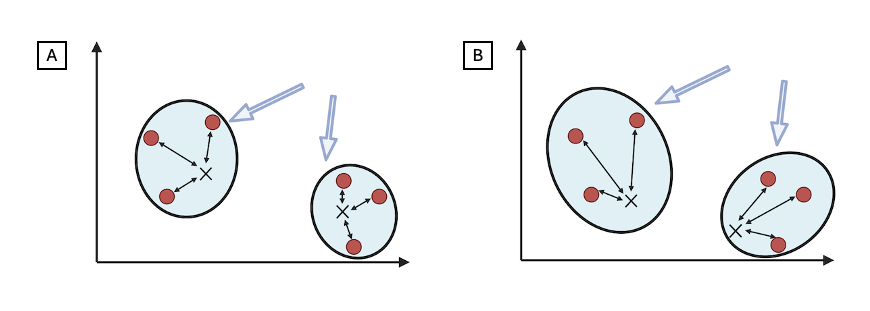

**Supplementary Fig.** 2. **The number of clusters to include is chosen prior to starting the analysis.** (A) Cluster centroids, represented by the x in the figure, are randomly chosen, and the Euclidean distance of each data point to the nearest centroid is calculated (Arthur & Vassilvitskii, n.d.). The total within sum of squares value is calculated for each version(Arthur & Vassilvitskii, n.d.). (B) Generating different centroid locations, the value of the total within sum of squares is compared, and the centroid location generating the smallest value for the total within sum of squares is selected.
